# Karanjin alters gene expression through ERα: a preliminary study

**DOI:** 10.1101/2024.03.11.584382

**Authors:** Gaurav Bhatt, Latha Rangan, Anil Mukund Limaye

**Affiliations:** Department of Biosciences and Bioengineering, Indian Institute of Technology Guwahati, Guwahati 781039, Assam, India

**Author notes:** Corresponding author Anil Mukund Limaye, Department of Biosciences and Bioengineering, Indian Institute of Technology Guwahati Guwahati 781039, Assam, India.

## Abstract

Karanjin is an abundant furanoflavonoid-constitutent of pongamia oil. Among several biological actions of karanjin, the antiproliferative effect of karanjin has gained traction in the recent years; raising speculations about its anticancer potential. In the backdrop of partial estrogen-like alteration of gene expression by karanjin in ERα positive MCF-7 breast cancer cells, we present preliminary evidences supporting of the role of ERα in karanjin-mediated effects.

## Introduction

Karanjin, a furanoflavonoid, is a major constituent of Pongamia seed oil, which has applications in *Ayurveda*, and *Siddha*. This molecule is known since 1925^1^. However, investigations into its biological actions through modern scientific methods is a relatively recent activity. Karanjin exerts diverse biological actions ^2,3^. Investigations in cell culture models have revealed cell cycle-inhibitory, or apoptosis-inducing effects, which highlight its anticancer potential^4–7^. Recently we published partial estrogen-like effects of karanjin in breast cancer cells^8^. Here, we communicate preliminary, yet significant, insights into the mechanism of its pro-estrogenic action that implicates the classical estrogen receptor-α (ERα).

## Materials and Methods

### Molecular docking protocol and validation

Eight wild-type agonist (17β-estradiol [E2])-bound, and two antagonist (4-hydroxy tamoxifen [OHT], or raloxifene [RAL])-bound three-dimensional crystal structures of ERα ligand binding domain (LBD) were retrieved from the Protein Data Bank^9^. The crystallographic ligands were extracted from the respective complexes, re-docked, and superimposed onto the co-crystallized complexes to validate the docking protocol using AutoDock 4.2.6^10^. For ERα, water molecules were removed, and polar hydrogen atoms were added. Gasteiger and Kollman charges were added. The 3D coordinates of the ligands E2, OHT, RAL, and karanjin were retrieved in Structure Data File (SDF) format from PubChem^11^. They were converted to PDB format using the online SMILES translator or Open Babel software^12^. All default torsions of the ligands were allowed to rotate during the docking run. A grid box 40 × 40 × 40 was prepared, encompassing the ligand binding pocket of ERα. Grid spacing of 0.375 Å was used to compute affinity, and electrostatic maps. Docking search was carried out using the Lamarckian genetic algorithm. All remaining parameters were set as default. The docked conformations were sorted according to the predicted binding energies, with the lowest energy conformation considered the most reliable. The conformations obtained were compared with the crystallographic ligand-bound ERα-LBD by determining the root-mean-squared deviation (RMSD). Upon redocking, E2, OHT, and RAL were found to occupy the ligand binding pocket of ERα, and the RMSD was close to 1 Å in all of the eight E2- and two OHT- or RAL-bound ERα-LBD structures (Supplementary data 1).

### Chemicals, reagents, and plasticware

Dulbecco’s Modified Eagle Medium (DMEM) with (Cat. No. AT007) or without phenol red (Cat. No. AT187), Dulbecco’s Phosphate Buffered Saline (DPBS, Cat. No. TS1006), trypsin-EDTA (Cat. No. TCL014), antibiotic solution (Cat. No. A001), fetal bovine serum (FBS, Cat. No. RM10432), and charcoal-stripped FBS (cs-FBS, Cat. No. RM10416), were purchased from HiMedia (Mumbai, India). PowerUp SYBR Green PCR Master Mix (Cat. No. A25743), High-Capacity cDNA Reverse Transcription Kit (Cat. No. 4368814), ERα siRNA (Cat No. 4392420), scrambled siRNA (Cat. No. AM4611), and Lipofectamine RNAiMAX (Cat. No. 13778075) were from Invitrogen (CA, USA). All other chemicals and buffers were purchased from Merck (Mumbai, India), Sigma (St Louis, MO, USA), or SRL (Mumbai, India). All cell culture plasticware were from Eppendorf (Hamburg, Germany). Karanjin standard was procured from Yucca Enterprises (CAS No. 521-88-0, Batch No. Yucca/KG/2019/04/21, Mumbai, India). Karanjin was dissolved in DMSO to a working stock concentration of 50 mM, and stored at -20ºC.

### Cell culture

MCF-7 cells were obtained from the National Centre for Cell Science (Pune, India). They were routinely cultured and expanded in phenol red-containing DMEM, which was supplemented with 10% heat-inactivated FBS, 100 units/mL penicillin, and 100 μg/mL streptomycin (M1 medium) under standard humidified conditions at 37°C and 5% CO_2_.

### Time course experiment

MCF-7 cells were seeded in 35 mm dishes, and incubated in M1 medium until 70% confluent. The cells were shifted to phenol red-free DMEM, supplemented with 10% heat-inactivated cs-FBS, 100 units/mL penicillin, and 100 μg/mL streptomycin (M2 medium) for 24 h. Then the cells were treated with vehicle (0.1% DMSO), or 10 µM karanjin for 0, 36, 48, and 72 h. For the 72 h treatment, the medium with karanjin was replenished once after 48 h. Following the completion of the experiment, the cells were processed for total protein isolation.

### Effect of ER_α_ knockdown on karanjin-modulated gene expression

2.5×10^5^ MCF-7 cells were seeded in six-well plates, and incubated for 24 h in M1 medium. Cells were transfected with ERα-specific, or scrambled (control) siRNA for 24 h using Lipofectamine RNAiMAX according to manufacturer’s instructions. Then, the cells were either treated with vehicle (0.1% DMSO), or 10 µM karanjin in M2 medium for 24 h. Thereafter, the cells were processed for total RNA isolation.

### Total RNA isolation, cDNA synthesis and RT-qPCR

Total RNA was isolated using an RNA extraction reagent made in-house according to Chomczynski and Sacchi^13^. RNA integrity was determined using agarose gel electrophoresis, and quantified using a BioSpectrometer® (Eppendorf, Hamburg, Germany). 2 µg of total RNA was reverse transcribed using the High-Capacity cDNA Reverse Transcription kit. The qPCR reactions were setup in AriaMX Real-time PCR system (Agilent, CA, USA) with 2 µl diluted cDNA (1:10), PowerUp™ SYBR™ Green PCR master mix, and gene-specific primers listed in Supplementary data 2. Cyclophilin A served as internal control.

### Western blotting

Total protein was extracted using the 1.5X Laemmli buffer^14^. Protein was quantified by TCA method. 30 µg of protein was fractionated on 10% denaturing SDS-PAGE gel, and transferred to a nitrocellulose membrane (Cat No. SF110B, HiMedia laboratories, Mumbai, India). The membrane was blocked for 2 h at room temperature with 1% gelatin (w/v) in tris-buffered saline containing 0.05% Tween 20 (TBST). The blots were probed for 2 h with anti-ERα (Cat. No. 8644S, Cell Signalling Technology, Massachusetts, USA), or histone H3 (Cat. No. BB-AB0055, BioBharati LifeScience Pvt. Ltd., Kolkata, India) antibodies followed by 1X TBST washes for 30 mins (6 washes of 5 mins each). The blots were incubated for 1 h in HRP-conjugated anti-rabbit secondary antibody (Cat. No. 7074S, Cell Signalling Technology, Massachusetts, USA), and washed again for 30 mins (6 washes of 5 mins each). The chemiluminescence signals were obtained with Clarity Western ECL Substrate (Bio-Rad, California, USA, Cat. No. 1705060) and captured with ChemiDoc XRS+ system (Bio-Rad, California, USA). Histone H3 served as an internal control.

### Statistical analysis

To study the effect of ERα knockdown on karanjin-modulated gene expression, the data were analysed by two-way ANOVA, after confirming homogeneity of variance using the Levene test, and normality, using the Shapiro-Wilk test. Two-group data were analysed by one-tailed t-tests. All statistical analyses were performed in R.

## Results and Discussion

The enrichment of estrogen-response-early genes in the karanjin-modulated transcriptome of the ERα-positive MCF-7 breast cancer cells^8^ instigated us to hypothesize ERα as a functional target. The availability of agonist- or antagonist-bound crystal structures of ERα ligand binding domain (LBD) facilitated *in silico* molecular docking experiments to explore karanjin-ERα interaction. The ERα ligand binding pocket comprises of 22 amino acids; with Leu 346, Ala 350, Leu 384, Leu 387, Phe 404, Val 418, Met 421, Ile 424, His 524, Glu 353, and Leu 525 identified as those involved in E2-ERα interaction^15^. Following validation of the molecular docking protocol as described in Materials and Methods, we docked karanjin to ERα-LBD extracted from eight wild-type agonist (E2)- or two antagonist (OHT or RAL)-bound structures. The docked conformations were visualized via LigPlot software v2.2. Analysis of the docked conformations revealed that karanjin also occupied the ERα ligand binding site characterized by interaction with Leu 346, Ala 350, Leu 387, Phe 404, Met 421, Ile 424, His 524, Glu 353, and Leu 525. However, unlike E2, karanjin failed to interact with two residues, namely Val 418 and Leu 383. Analysis of karanjin interaction in all eight E2-bound crystal structures suggested that the furan ring-oxygen of karanjin has polar interaction with His 524. Additionally, the remainder of the karanjin molecule participated in a number of hydrophobic contacts with residues in helices 3, 6, 11, and 12 (Supplementary data 3).

The agonist and antagonist ligands bind to the same site within the core of the LBD of ERα, but exhibit different binding modes. The helix 12 conformation of the ERα LBD differs; depending upon the nature of the ligand. There is a change in helix 12 conformation due to the absence of a hydrogen bond with His 524, when OHT is bound^15^. In contrast, hydrogen bonding occurs with His 524 in the agonist E2-bound conformation. Docking conformations of karanjin with E2-bound (2YJA), and OHT-bound (3ERT) ERα structures showed the interaction of karanjin with His 524 in the former, but not in the latter (Fig. 1A). This suggests that karanjin may be a potential ERα agonist, which is possibly reflected in the partial E2-like effects of karanjin described earlier^8^.

**Fig. 1.**
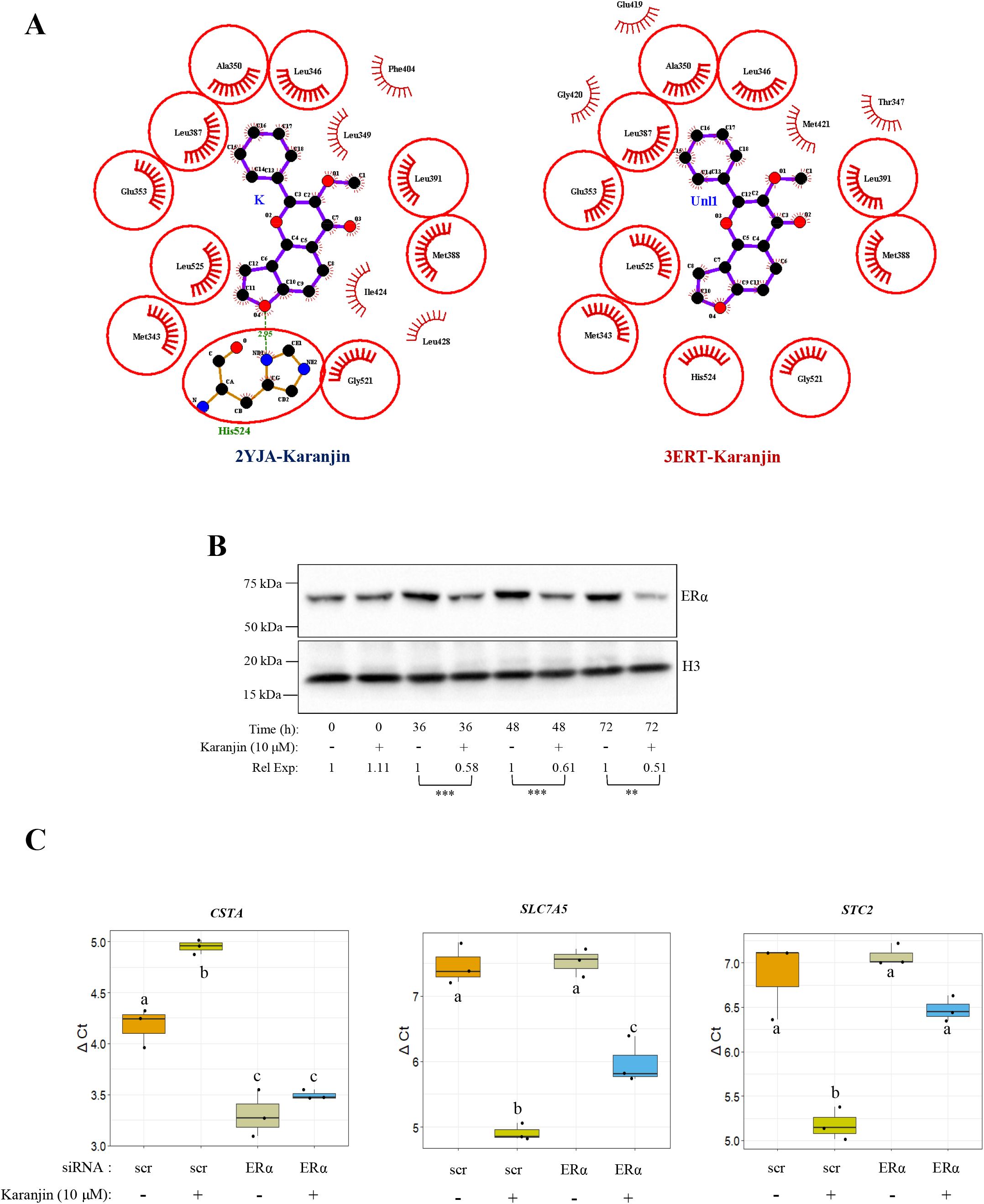
*In-silico*, and experimental evidences indicating ER_α_ as a mediator of karanjin action. **A**. ERα structures extracted from the E2-bound (2YJA) or OHT-bound (3ERT) crystallographic complexes were docked with karanjin as described in Materials and Methods. The docked conformations were visualized via Ligplot+-derived 2D interaction diagrams. Karanjin is tethered in the centre surrounded by the contacting amino acid residues (red color) that are described as circles and ellipses. In 2YJA-karanjin complex, the hydrogen bonded residue His 524 is highlighted, and green dotted line represents hydrogen bond formation with furan ring oxygen of karanjin in E2-bound structure (left top panel), but not in 3ERT-karanjin structure (right top panel). **B**. Karanjin-mediated reduction in ERα protein expression in MCF-7 cells. Cells were treated with 10 µM karanjin for the indicated period of time. At each time point, the untreated cells served as control. Total protein extracted in 1.5X Laemmli buffer was subjected to western blotting analysis using the ERα-specific antibody. Histone H3 served as an internal control, which was probed with H3-specific antibody. Chemiluminescence signals were imaged in the ChemiDoc XRS+ system. The images were processed in ImageJ v1.53t. The background-subtracted band intensity for ERα, which was normalized against that obtained for Histone-H3 served as a measure of ERα protein expression. For each time point, ERα expression in the control sample was set to 1, and that obtained for the karanjin-treated sample was expressed relative to the control. The mean relative expression (Rel Exp) of ERα ± sd (n = 5) for each time-point is shown below. For each time point, the data were analysed by a one-tailed t-test. Significant results are indicated by asterisks. (^*^ p < 0.05, ^**^ p < 0.01, ^***^p < 0.001). **C**. ERα knockdown affects modulation of gene expression by karanjin. MCF-7 cells pre-treated with scrambled (scr) or ERα-specific siRNA were treated with vehicle or 10 µM karanjin for a period of 24 h. Total RNA was extracted from MCF-7 cells, and the expression of the indicated genes was analysed by RT-qPCR. The experiment was done with three replicate dishes for each experimental group. For each sample the average Ct value for CSTA, and the internal control CycA was determined, and the difference, which is referred to as ΔCt, was considered as normalized expression. As a result, higher ΔCt reflects lower normalized expression for a given gene, and *vice versa*. For each gene, the data were analysed by two-way ANOVA followed by TukeyHSD. The statistical difference between the pairs of treatment is denoted by a, b, and c.

The *in-silico* data motivated us to investigate the role of ERα in karanjin-mediated effects. Treatment of MCF-7 cells with 10 µM karanjin decreased the steady-state levels of ERα protein (Fig. 1B), which recapitulates the known effect of E2^16^. CSTA, STC2, and SLC7A5 are E2-modulated genes^17–19^. Previously, we showed that these genes are modulated by karanjin^8^. To test whether karanjin modulated CSTA, STC2, or SLC7A5 via ERα, MCF-7 cells were transfected with scrambled or ERα-specific siRNA, and treated with vehicle or karanjin, and their mRNA expression levels were determined by RT-qPCR. For each sample, the threshold cycle number was determined for the gene of interest and the normalizing control (Cyc A). The difference (ΔCt) was determined as a measure of the normalized expression in each sample. Thus, larger ΔCt reflected lower expression. The data were analysed by two-way ANOVA, to study the main effects of karanjin, or ERα knockdown, or their interaction, if any. Figure 1C, is a graphical representation of the data. There were significant main effects of karanjin treatment (p < 0.001 for CSTA, p ≈ 0 for SLC7A5, p ≈ 0 for STC2), or ERα knockdown (p ≈ 0 for CSTA, p = 0.0069 for SLC7A5, p < 0.001 for STC2), suggesting that both independently altered the expression of these genes. However, the interaction between karanjin, and ERα knockdown was also significant (p = 0.011 for CSTA, p = 0.011 for SLC7A5, and p = 0.0062 for STC2), suggesting that the effect of karanjin on the expression of these genes was significantly affected by ERα knockdown. The results of the two-way ANOVA including the p values resulting from pair-wise comparison of the ΔCt values among the treatment groups are presented as Supplementary data 4. Karanjin treatment significantly decreased the normalized expression of CSTA mRNA in scrambled siRNA treated MCF-7 cells, not in those treated with ERα specific siRNA (Fig 1C, left panel). On the other hand, karanjin treatment significantly induced the expression of SLC7A5 and STC2 in MCF-7 cells treated with scrambled siRNA. However, ERα siRNA partially or completely blocked karanjin mediated induction in SLC7A5 (Fig 3C, middle panel), or STC mRNA (Fig 3C right panel), respectively. Overall, these results provide compelling evidences to implicate ERα, at least in part, as a mediator of karanjin effects.

Flavonoids are known for their estrogenic activity^20^. Given the flavonoid nature of karanjin, our results, though not surprising, presents a fresh perspective on the mechanism of karanjin action that contrasts the antiproliferative effects described by others. In-depth molecular investigations into the estrogen receptor modulatory action of karanjin, or its tailored derivatives, is worthwhile; for it provides a compelling lead for development of novel endocrine therapies.

## Acknowledgments

The work was supported by financial assistance from Science and Engineering Research Board, Department of Science and Technology, Govt. of India (File No. CRG/2020/002109). We acknowledge the support from the DBT funded Bioinformatics Facility, and infrastructural support from IIT Guwahati.

## References

1. Limaye, D. B. Karanjin part I: a crystalline constituent of the oil from Pongamia glabra. Proc. 12th Indian Acad. Sci. Congr. India. 118, 118–125 (1925).

2. Mohd Noor, A. A. et al. Molecules of Interest – Karanjin – A Review. Pharmacogn. J. 12, 938–945 (2020).

3. Singh, A. et al. Karanjin. Phytochemistry 183, (2021).

4. Guo, J.-R., Chen, Q.-Q., Lam, C. W.-K. & Zhang, W. Effects of karanjin on cell cycle arrest and apoptosis in human A549, HepG2 and HL-60 cancer cells. Biol. Res. 48, 40 (2015).

5. Roy, R., Mandal, S., Chakrabarti, J., Saha, P. & Panda, C. K. Downregulation of Hyaluronic acid-CD44 signaling pathway in cervical cancer cell by natural polyphenols Plumbagin, Pongapin and Karanjin. Mol. Cell. Biochem. 476, 3701–3709 (2021).

6. Roy, R. et al. Pongapin and Karanjin, furanoflavanoids of Pongamia pinnata, induce G2/M arrest and apoptosis in cervical cancer cells by differential reactive oxygen species modulation, DNA damage, and nuclear factor kappa-light-chain-enhancer of activated B cell signal. Phytother. Res. 33, 1084–1094 (2019).

7. Yu, J., Yang, H., Lv, C. & Dai, X. The cytotoxicity of karanjin toward breast cancer cells is involved in the PI3K/Akt signaling pathway. Drug Dev. Res. 83, 1673–1682 (2022).

8. Bhatt, G., Gupta, A., Rangan, L. & Mukund Limaye, A. Global transcriptome analysis reveals partial estrogen-like effects of karanjin in MCF-7 breast cancer cells. Gene 830, 146507 (2022).

9. Berman, H. M. et al. The Protein Data Bank. Nucleic Acids Res. 28, 235–42 (2000).

10. Forli, S. et al. Computational protein-ligand docking and virtual drug screening with the AutoDock suite. Nat. Protoc. 11, 905–19 (2016).

11. Kim, S. et al. PubChem in 2021: new data content and improved web interfaces. Nucleic Acids Res. 49, D1388–D1395 (2021).

12. O’Boyle, N. M. et al. Open Babel: An open chemical toolbox. J. Cheminform. 3, 33 (2011).

13. Chomczynski, P. & Sacchi, N. Single-step method of RNA isolation by acid guanidinium thiocyanate-phenol-chloroform extraction. Anal. Biochem. 162, 156–9 (1987).

14. Laemmli, U. K. Cleavage of structural proteins during the assembly of the head of bacteriophage T4. Nature 227, 680–5 (1970).

15. Brzozowski, A. M. et al. Molecular basis of agonism and antagonism in the oestrogen receptor. Nature 389, 753–8 (1997).

16. Reid, G. et al. Cyclic, proteasome-mediated turnover of unliganded and liganded ERalpha on responsive promoters is an integral feature of estrogen signaling. Mol. Cell 11, 695–707 (2003).

17. John Mary, D. J. S., Manjegowda, M. C., Kumar, A., Dutta, S. & Limaye, A. M. The role of cystatin A in breast cancer and its functional link with ERα. J. Genet. Genomics 44, 593–597 (2017).

18. Charpentier, A. H. et al. Effects of estrogen on global gene expression: identification of novel targets of estrogen action. Cancer Res. 60, 5977–83 (2000).

19. Bouras, T. et al. Stanniocalcin 2 is an estrogen-responsive gene coexpressed with the estrogen receptor in human breast cancer. Cancer Res. 62, 1289–95 (2002).

20. Zand, R. S., Jenkins, D. J. & Diamandis, E. P. Steroid hormone activity of flavonoids and related compounds. Breast Cancer Res. Treat. 62, 35–49 (2000).

