## Supplementary data 1 for "Karanjin alters gene expression through ERα: a preliminary study"

**Supplementary data 1:** List of estrogen-bound PDB structures of wild type ERα LBD.

| **PDB** | **Resolution** | **Chains** | **Length (amino acids)** | **Ligand** | **RMSD** |
| --- | --- | --- | --- | --- | --- |
| 1A52 | 2.8 | A/B | 297-554 | E2 | 0.6 A |
| 1ERE | 3.1 | A/B/C/D/E/F | 301-553 | E2 | 0.6 A |
| 1G50 | 2.9 | A/B/C | 304-550 | E2 | 0.5 A |
| 1GWR | 2.4 | A/B | 305-549 | E2 | 0.7 A |
| 1QKU | 3.2 | A/B/C | 301-550 | E2 | 0.5 A |
| 2YJA | 1.82 | B | 299-551 | E2 | 0.6 A |
| 5GS4 | 2.4 | A | 305-547 | E2 | 0.4 A |
| 5GTR | 2.8 | A | 305-547 | E2 | 0.9 A |
| 3ERT | 1.9 | A | 299-551 | OHT | 1.1 A |
| 1GWQ | 2.45 | A/B | 302-549 | RAL | 0.9 A |
