## Supplementary data 2 for "Karanjin alters gene expression through ERα: a preliminary study"

**Supplementary table 2:** List of primers.

| **Primer name** | **Sequence (5’-3’)** | **Amplicon length** | **Annealing temp.** |
| --- | --- | --- | --- |
|  |  | **(bp)** | **(°C)** |
| CSTA-F | ATCTGAGGCCAAACCCGCC | 275 | 60 |
| CSTA-R | AGCCCGTCAGCTCGTCATC |  |  |
| CycA-F | GGGCCGCGTCTCCTTTGAGC | 158 | 60 |
| CycA-R | GGCGTGTGAAGTCACCACCC |  |  |
| SLC7A5-F | GTGGACTTCGGGAACTATCACC | 126 | 60 |
| SLC7A5-R | GGACCCCACGAAGAAGAGC |  |  |
| STC2-F | ATGCTACCTCAAGCACGACC | 129 | 60 |
| STC2-R | CAGGTCAGCAGCAAGTTCAC |  |  |

Note: F and R indicate sense and antisense primers, respectively.
