## Supplementary data 3 for "Karanjin alters gene expression through ERα: a preliminary study"

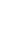

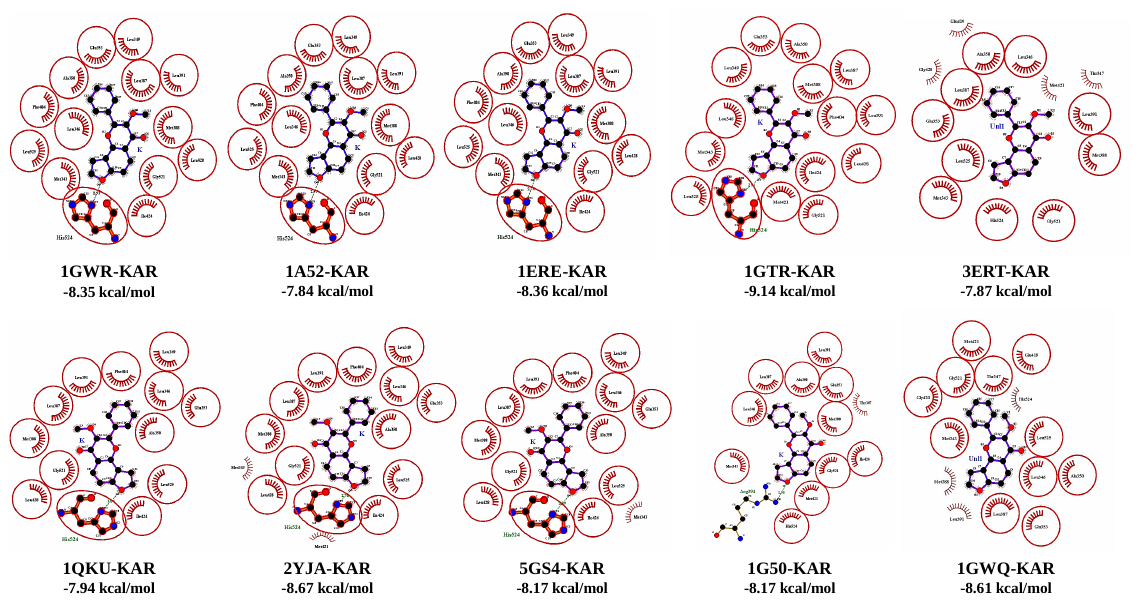
**Supplementary data 3: Karanjin-docked ERα LBD crystal structures visualized in LigPlot+**. The docking was performed as described in Materials and Methods provided as Supplementary data 1. 1GWR, 1A52, 1ERE, 1GTR, 1QKU, 2YJA, 5GS4, 1G50 are E2-bound crystallographic PDB structures docked with karanjin (KAR). Similarly, 3ERT and 1GWQ are antagonist-bound crystallographic structures docked with KAR. The values beneath each complex are the predicted binding energy in Kcal/mol. The contacting amino acids (red color) are described as circles and ellipses. The hydrogen-bonded residue, His 524 is highlighted, and the green dotted line represents hydrogen bond formation with furan ring oxygen of karanjin in E2-bound structures, but not in antagonist bound structures.
